# fastQpick: scalable bootstrap and subsampling of FASTQ reads

**DOI:** 10.64898/2026.06.23.734068

**Authors:** Joseph Rich, Lior Pachter

## Abstract

Summary: fastQpick is a command-line tool and Python library for sampling FASTQ reads with replacement. Sampling with replacement turns a single FASTQ file into an arbitrary number of bootstrap replicates, which enables uncertainty quantification and statistical analysis at the level of raw reads. This process answers questions such as how much an abundance estimate would change if the library were resequenced, or whether a low-abundance call is robust to the particular reads that were sequenced. fastQpick works efficiently on large libraries by streaming files in two passes by default: first to count reads and create a hash-based counter, and then to write the sample. It generates a full-size bootstrap replicate of a 500-million-read library in under 30 minutes with 9.4 GB of peak memory, with a low-memory mode that reduces the peak to 1.4 GB. A single-pass mode draws samples in a single read through the file, using *O*(1) working memory and producing an output size that is exact in expectation but not fixed. In a real yeast RNA-seq experiment, bootstrap replicates generated by fastQpick recover the sampling uncertainty of transcript abundance estimates, matching the analytic multinomial standard errors to within a few percent.

**Availability and implementation:** fastQpick is open source and freely available under the MIT license on GitHub at https://github.com/pachterlab/fastQpick and on PyPI (`pip install fastQpick`).

## Introduction

A result computed from a sequencing library such as a transcript’s abundance, a variant’s allele frequency, or a taxon’s relative abundance is an estimate from a single random sample of the molecules in that library. Yet this result is typically reported as a point value with no measure of how much it would change if the library were sequenced again. Quantifying this uncertainty can be difficult in the absence of biological replicates, as many statistics of interest have no closed-form standard error. The nonparametric bootstrap (Efron, 1979) addresses both cases by resampling the observed data with replacement to the same size as the original data set and recomputing the statistic, repeating this many times to approximate its sampling distribution. Applied at the level of raw reads, it propagates sequencing sampling uncertainty through any downstream pipeline and yields an error bar for any statistic. For example, fastQpick can assess whether a low-abundance variant or transcript call is robust to the particular reads that were sequenced, place confidence intervals on quantities that lack a closed-form standard error such as ecological diversity indices or abundances derived from probabilistic assignment of multi-mapping reads, and compute *p*-values for whether a biological effect exceeds the variability introduced by library preparation and sequencing.

The value of resampling reads with replacement directly from FASTQ files has been established in the context of reproducibility assessment. Saremi et al. (2020) introduced bootstrap sampling of reads from FASTQ files to measure the reproducibility of virus metagenomics analyses, showing that bootstrap replicates reveal which viral identifications are stable and which are driven by a small number of reads. Saremi et al. (2022) compared strategies for generating artificial replicates in RNA-seq experiments and found read-level resampling to be an effective way to produce replicates for downstream statistical analysis.

Most general-purpose read sampling, by contrast, is performed without replacement, as a common preprocessing step in DNA-seq and RNA-seq workflows. It is used to equalize sequencing depth across libraries before comparison, to produce smaller inputs for development and testing, and to study how sensitive a result is to the particular reads that were sequenced. The standard tools for this task, such as `seqtk sample` (Li, 2018), `seqkit sample` (Shen et al., 2016), and the subsampling mode of `fastp` (Chen et al., 2018), draw a subset of reads without replacement.

Sampling without replacement is a relatively straightforward task. A tool such as `seqtk` (Li, 2018) can draw an exact number of reads in a single pass using reservoir sampling, which holds the *k* reads to be kept and updates this reservoir as each read streams past, so the total read count *n* never needs to be known in advance. Sampling with replacement is a more challenging task, because each output read is drawn uniformly from the entire file, so the total read count *n* must be known before any read can be sampled. This requires one pass to count the reads and a second pass to write the sample, since the alternative of holding the entire file in memory is infeasible for libraries of hundreds of millions of reads. The two-pass approach trades the cost of reading the file a second time for an exact output size at memory proportional to the read count rather than the read content. A single-pass alternative avoids the counting pass by drawing each read’s number of output copies as it streams by (Poisson with replacement, Bernoulli without), which uses constant memory and a single read through the file but yields an output size that is random and exact only in expectation.

fastQpick enables sampling of FASTQ reads with (or without) replacement at any fraction of the input. The default mode balances exactness, memory, and time efficiency, with options to optimize any two of these at the cost of the third. It scales to libraries of hundreds of millions of reads, running bootstrapping in under 30 minutes for a 500-million-read library.

## Methods

### User interface

fastQpick is invoked either from the command line or from Python, and both entry points call the same underlying routine. The only required argument is the list of input files. By default, sampling is performed with replacement (`without replacement=False`) and creates output files that are the same size as the input (`fraction=1.0`), which is the standard bootstrap procedure. `fraction<1.0` performs subsampling, and `fraction>1.0` performs oversampling with replacement. Random seeds can be specified with the `seed` argument to make the sampling reproducible, and the `num samples` argument generates multiple independent replicates in a single invocation. Additional arguments exist to control the output directory, gzip compression, file grouping (e.g., paired-end files, or I1/R1/R2 triples), low-memory mode, single-pass mode, and unique header names. Type validation and conversion are handled by the Python package `pydantic` (Colvin et al., 2026).

### Data structures

Let *n* be the number of reads in a file and let *m* = ⌊*f* · *n*⌋ be the number of reads to sample for a requested fraction *f* . Processing proceeds in two passes over the file, both streamed through `pyfastx` so that read records are never held in memory all at once.

The first pass counts the reads in each file. For grouped files, only the first member is counted and the count is reused for the rest of the group, which requires that grouped files contain equal numbers of reads. The count can also be supplied by the caller to skip this pass entirely.

Given the count, fastQpick draws *m* read indices in [0, *n*) and converts them into an occurrence vector whose *i*-th entry records how many times read *i* is to be written. Two data-structure choices keep this step memory-efficient. The first is an adaptive occurrence structure: when the fraction *f* is small, occurrences are stored in a hash-based counter keyed only by the sampled indices, using memory proportional to the number of distinct reads selected. Otherwise, they are stored in a dense integer array of length *n* built by a counting (bincount) operation. The threshold is set where the per-entry overhead of the counter, on the order of one hundred bytes, equals the one byte per read of the dense array, which places it near *f* = 0.01; the two representations are interchangeable downstream because both map a read index to its occurrence count. Because each count is distributed as Poisson(*f* ) with replacement and is at most one without replacement, this width is a single byte in essentially all practical cases, so the dense array costs about one byte per read. The index container is sized separately to the smallest type that can represent *n*. The peak memory of this step is set not by the persistent occurrence vector, but rather transient inputs to the bincount operation, namely the length-*m* array of sampled indices together with the length-*n* counting buffer it produces. Both of these objects are released once the occurrence vector has been formed.

A low-memory mode replaces the materialized index array with lazy generators built on the standard-library `random.sample` and `random.choice`, accumulating directly into the occurrence array. This avoids both the array of *m* candidate indices and the length-*n* counting temporary, at the cost of a small increase in preprocessing time. The two modes use similar memory during the writing pass, since both write from the same one-byte-per-read occurrence array, but the default mode has a higher transient peak while the occurrence array is built.

The second pass streams the file once more and writes each read as many times as its occurrence count, accumulating output in a buffer that is flushed in batches to amortize the cost of writing. For grouped files, the occurrence vector computed from the first member is reused for every member, so all passes after the first reuse the same sampling decision.

### Single-pass sampling

The counting pass exists only to determine *n*, which the default sampler needs because drawing exactly *m* = ⌊*f* · *n*⌋ reads is a constraint on the file as a whole. For a uniform random sample, the following three properties cannot be attained together: a single pass over the file, *O*(1) working memory, and an exactly determined output size. The default sampler attains the last two at the cost of an extra pass. An optional single-pass mode attains the first two by relaxing the output size to be exact only in expectation.

In single-pass mode the output multiplicity *c*_*i*_ of read *i* is drawn independently for each read as the file streams by, so the file length is never required. With replacement, *c*_*i*_ ∼ Poisson(*f* ); without replacement, *c*_*i*_ ∼ Bernoulli(*f* ), so a read appears at most once. The Poisson choice is the Poissonized form of with-replacement sampling: in an exact draw of *m* reads with replacement the perread counts are multinomial, and the marginal count of any one read is Binomial(*m*, 1*/n*), which converges to Poisson(*m/n*) = Poisson(*f* ) as *n* grows. Poissonization makes the per-read counts independent, which removes the dependence on the total *m* and hence on *n*.

Both rules satisfy E[*c*_*i*_] = *f* for every read, independent of its position *i*. Therefore, the expected output size is 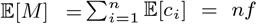, where 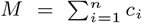 is the random variable for the number of reads sampled. This contrasts with reservoir sampling (Vitter, 1985), the other single-pass alternative, whose acceptance probability depends on read position and which requires memory proportional to the output size. With replacement, *M* ∼ Poisson(*fn*), giving standard deviation 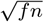; without replacement, *M* ∼ Binomial(*n, f* ), giving standard deviation 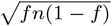. In both cases, the relative standard deviation is at most 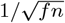, which is negligible at scale: for *f* = 0.1 and *n* = 10^8^, it is approximately 3 × 10^−4^. For grouped files, synchronization is maintained by using the same random seed for each file in a group.

### Complexity

In the default mode, the first pass (for counting total reads) works in *O*(*n*) time and *O*(1) memory. Drawing indices and building the occurrence structure take *O*(*n* + *m*) time. When *f* < 0.01, the occurrence structure takes *O*(*m*) memory (for the hash-based counter), else it takes *O*(*n*) memory (for the dense array). The second pass (for writing the sample) also works in *O*(*n* + *m*) time, since it visits all *n* input reads and emits *m* output reads. The overall time and space complexity is therefore *O*(*n* + *m*), which is *O*(*n*) for any fixed fraction.

The single-pass mode runs in *O*(*n*+*M* ) time, dominated by reading the *n* input reads once and writing the *M* output reads. Its peak memory is *O*(1), as no occurrence structure is built.

### Runtime and Memory Benchmarks

The runtime and memory usage of the three sampling modes were benchmarked on a 500-million-read RNA-seq library (Table 1). fastQpick generates a full-size bootstrap replicate (`fraction=1.0`, `without replacement=False`) in about 26 minutes in the default mode, about 56 minutes in low-memory mode, and about 35 minutes in single-pass mode, with runtime dominated by input and output rather than by the sampling itself. Peak resident memory is about 9.4 GB (about 20 bytes per read) in the default mode, about 1.4 GB in low-memory mode (about 3 bytes per read), and about 0.11 GB in single-pass mode. At `fraction=1.0` the default mode reads the file once into the page cache and re-reads it from cache during the writing pass, so on a machine whose memory holds the entire file it is faster in wall-clock time than the single-pass mode, which reads and writes concurrently and contends for the disk even though it moves less total data. This is demonstrated by how, at `fraction=0.01`, the single-pass mode is faster (4 minutes) than the default mode (8 minutes).

**Table 1.**
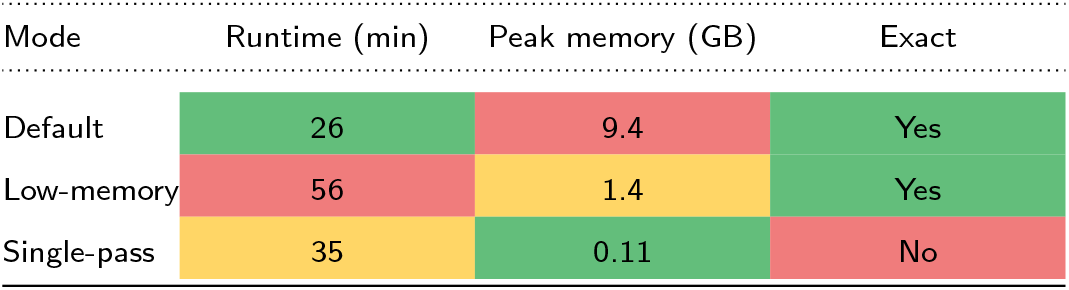
Performance of the three fastQpick sampling modes on a full-size bootstrap of a 500-million-read library. Green = strong; yellow = moderate; red = weak.

## Application

Read-level bootstrapping is demonstrated on a paired-end *Saccharomyces cerevisiae* RNA-seq library with 5,725,730 read pairs (Nookaew et al., 2012). The reads are quantified against the Ensembl R64-1-1 transcriptome (6612 transcripts) with kallisto (Bray et al., 2016), of which 5,190,446 reads (90.7%) pseudoalign. From the single library, fastQpick generates *B* = 200 full-size bootstrap replicates. Each replicate is re-quantified with kallisto, which yields a bootstrap distribution of the estimated proportion 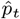 of pseudoaligned reads assigned to each transcript *t*.

The read-level bootstrap recovers the sampling uncertainty of the abundance estimates directly from the resampled reads. For a proportion estimated from *N* reads, the analytic multinomial sampling error is 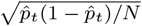, with *N* the number of pseudoaligned reads. Across the 4963 transcripts with at least 100 estimated counts, the standardized bootstrap deviations pooled across transcripts follow the standard normal distribution (Figure 1A), and the bootstrap standard error of 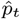 matches this analytic error closely (median relative difference 3.6%; Figure 1B). The bootstrap standard errors are on average 1.8% larger than the analytic multinomial errors, consistent with a small additional contribution from the probabilistic assignment of reads compatible with more than one transcript, which the multinomial term does not capture.

**Figure 1.**
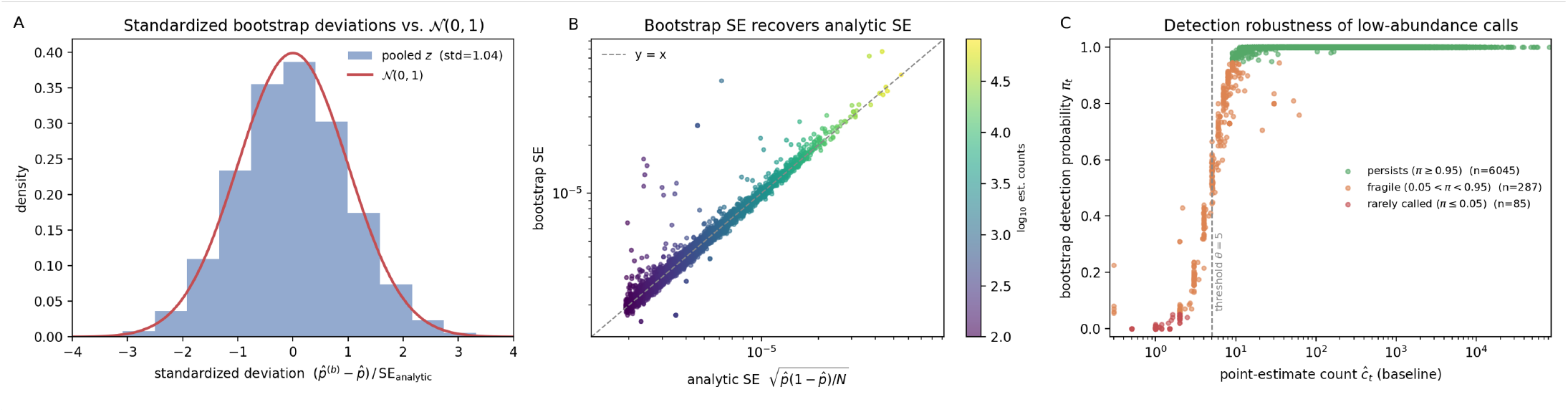
Read-level bootstrapping with fastQpick on a yeast RNA-seq library quantified with kallisto. (A) Standardized bootstrap deviations of the estimated transcript proportions pooled across all 4963 expressed transcripts and the 200 replicates. Red curve = standard normal density. (B) Bootstrap standard error of each transcript proportion against the analytic multinomial standard error, colored by estimated read count. Dashed line = *y* = *x*. (C) Bootstrap detection robustness of low-abundance transcripts. Fraction of bootstrap replicates in which each transcript’s estimated count reaches a detection threshold *θ* = 5 (dashed line) against the point estimate of its count from the original library, colored by detection fraction thresholds (green *π*_*t*_ ≥ 0.95, orange 0.05 < *π*_*t*_ < 0.95, red *π*_*t*_ ≤ 0.05).

The bootstrap also answers questions that have no closed-form solution, such as whether a low-abundance transcript call is robust to the particular reads that were sequenced. Detection is a binary outcome (presence or absence), so the per-transcript standard error of Figure 1B does not describe it. The bootstrap instead assigns each transcript a detection probability *π*_*t*_, the fraction of the *B* replicates in which its estimated count reaches a detection threshold *θ* = 5 (Figure 1C). Of the transcripts observed in the original library, 6045 persist with *π*_*t*_ ≥ 0.95, while 287 are fragile (0.05 < *π*_*t*_ < 0.95), i.e., their detection depends on the particular reads sequenced. Among these, 187 are called at baseline (*ĉ*_*t*_ ≥ *θ*) yet vanish in some replicates, and 100 are absent at baseline yet appear in others. The point estimate alone reports each of these transcripts as a definite presence or absence, whereas the read-level bootstrap flags which calls are driven by a handful of reads, the same robustness distinction that motivates resampling reads for reproducibility assessment (Saremi et al., 2020).

## Conclusion

fastQpick is a fast, memory-efficient tool for sampling FASTQ reads with replacement.

Its with-replacement mode, uncommon among general-purpose subsampling tools, makes read-level bootstrapping straightforward and brings nonparametric uncertainty quantification to the level of raw sequencing data. The two-pass streaming design, adaptive occurrence structures, minimal integer widths, and synchronized handling of grouped files allow the tool to scale to libraries of hundreds of millions of reads while keeping the interface to a single required argument.

## Data availability

No new data were generated for this study. The yeast RNA-seq library analyzed in the application is publicly available from the Sequence Read Archive under accession SRR453566 (Nookaew et al., 2012).

## Code availability

fastQpick and the notebook reproducing the application example are available on GitHub at https://github.com/pachterlab/fastQpick and on PyPI (`pip install fastQpick`).

## Author contributions statement

Joseph Rich (Conceptualization [lead], Data curation [lead], Formal analysis [lead], Investigation [lead], Methodology [lead], Software [lead], Validation [lead], Visualization [lead], Writing—original draft [lead], Writing—review & editing [lead]), and Lior Pachter (Conceptualization [supporting], Funding acquisition [lead], Methodology [supporting], Project administration [lead], Resources [lead], Supervision [lead], Writing—review & editing [lead]).

